# Different temporal windows for contextual fear memory destabilisation in the amygdala and hippocampus

**DOI:** 10.1101/434407

**Authors:** Jonathan L. C. Lee, Felippe E. Amorim, Lindsey F. Cassini, Olavo B. Amaral

## Abstract

Reconsolidation is a process in which re-exposure to a reminder causes a previously acquired memory to undergo a process of destabilisation followed by subsequent restabilisation. Different molecular mechanisms have been postulated for destabilisation in the amygdala and hippocampus, including CB1 receptor activation, protein degradation and AMPA receptor exchange; however, most of the amygdala studies have used pre-re-exposure interventions, while those in the hippocampus have performed them after re-exposure. To test whether the temporal window for destabilisation is similar across both structures, we trained Lister Hooded rats in a contextual fear conditioning task, and 1 day later performed memory re-exposure followed by injection of either the NMDA antagonist MK-801 (0.1 mg/kg) or saline in order to block reconsolidation. In parallel, we also performed local injections of either the CB1 antagonist SR141716A or its vehicle in the hippocampus or in the amygdala, either immediately before or immediately after reactivation. Infusion of SR141716A in the hippocampus prevented the reconsolidation-blocking effect of MK-801 when performed after re-exposure, but not before it. In the amygdala, meanwhile, pre-reexposure infusions of SR141716A impaired reconsolidation blockade by MK-801, although the time-dependency of this effect was not as clear as in the hippocampus. Our results suggest the temporal windows for CB1-receptor-mediated memory destabilisation during reconsolidation vary between brain structures. Whether this reflects different time windows for engagement of these structures or different roles played by CB1 receptors in destabilisation across structures remains an open question for future studies.

---

Memory reconsolidation is a core process in the maintenance and updating of long-term memories (1). Re-exposure to reminders reactivates previously learned memories, which may lead not only to their behavioural expression but also to reconsolidation (2). As reconsolidation depends upon neurochemical and cellular mechanisms of synaptic plasticity, such as NMDA receptor activation (3) and protein synthesis (4), pharmacological treatment around the time of memory reactivation can disrupt reconsolidation and result in subsequent amnesia.

Importantly, memory reactivation does not necessarily trigger reconsolidation (5). Instead there is a necessity for synaptic destabilisation, which has been shown to be dissociable from behavioural expression of the memory (6–8). Activation of GluN2B-containing NMDA receptors in the basolateral amygdala (BLA) was required for the destabilisation of auditory cued fear memories, whereas antagonism of AMPA or GluN2A-containing NMDA receptors selectively disrupted expression of the cued fear (6, 7).

Beyond GluN2B-containing NMDA receptors, little is known about the neurochemical mechanisms of memory destabilisation in the BLA, although AMPA receptor subunit exchange (9), synaptic protein degradation (10, 11), calcineurin (12), nitric oxide (13) and acidosis (14) have all been implicated (15). The requirement of AMPA receptor endocytosis and proteasome-mediated protein degradation recapitulate previous findings concerning hippocampal contextual fear memory destabilisation (16, 17), suggesting that there are common mechanisms of memory destabilisation across neural loci. Moreover, given that contextual fear conditioning relies critically upon both the dorsal hippocampus (DH) and the BLA (18, 19), synaptic destabilisation is likely to occur in both loci upon memory reactivation. Indeed, protein degradation is required not only in the DH but also in the BLA for the destabilisation of contextual fear memories (10).

The temporal requirement for destabilisation mechanisms in both structures, however, has not been extensively explored. Initially, it was shown that infusion of the GluN2B receptor antagonist ifenprodil into the BLA successfully disrupted memory destabilisation only when performed pre-reactivation (6). On the other hand, the observations that post-reactivation intra-DH infusions of a LVGCC channel blocker or cannabinoid CB1 receptor antagonist prevented the destabilisation of contextual fear memory (20) and that intracellular targeting of the proteasome after reactivation prevented destabilisation both in the hippocampus and amygdala (10, 16) suggest that the temporal dynamics of destabilisation might differ between these structures. However, this comparison is complicated by the fact that these studies differed not only in the locus of drug infusion, but also in the neurochemical target and the type of fear conditioning assessed (i.e. cued vs contextual).

To address this question, we aimed to directly compare the effects of pre-and post-reactivation drug infusion in the BLA and DH on the destabilisation of contextual fear memory. As our established reconsolidation-blocking drug, MK-801 (4) is a non-competitive NMDA receptor antagonist, we chose not to target GluN2B-related mechanisms, and instead focussed on CB1 receptor involvement in destabilisation (21, 22).

## Methods

### Subjects

143 male Lister Hooded rats (275-325 g at the time of surgical preparation), were housed in quads under a 12 h light/dark cycle (lights on at 0700) at 21°C with food and water provided ad libitum except during the behavioural sessions. Standard cages contained aspen chip bedding and environmental enrichment was available in the form of a Plexiglass tunnel. Experiments took place in a behavioural laboratory between 0830 and 1500. At the end of the experiment, animals were humanely killed via a rising concentration of CO2; death was confirmed by cervical dislocation. All procedures were approved by a local ethical review committee and conducted in accordance to the United Kingdom Animals (Scientific Procedures) Act 1986, Amendment Regulations 2012 (PPL P8B15DC34).

### Surgical preparation

All rats were implanted with chronic indwelling stainless steel cannulae (Coopers Needleworks, UK) under isoflurane anaesthesia and aseptic conditions according to our established procedures (23). 57 rats had cannulae targeting the DH (24) and the remaining 86 rats had cannulae targeting the basolateral amygdala (25). Cannula placements were verified by Nissl-staining of sectioned drop-perfused brains. Rats were included in the data analysis if there was histological evidence (glial scars) for the injector tip being located within the DH (including CA1, DG & CA3) or BLA (including all subregions of the Lateral Amygdaloid Nucleus and Basolateral Amygdaloid Nucleus). 9 rats were excluded from the DH groups, all on the basis of histological assessment. 37 rats were excluded from the BLA groups; 15 were unable to be infused bilaterally, and 22 were excluded on histological basis.

### Drugs

MK-801 (Abcam, UK) was dissolved in sterile saline to a concentration of 0.1 mg/ml and was administered i.p. at a dose of 0.1 mg/kg (21). SR141716A (Tocris, UK) was dissolved in a vehicle solution containing 3 drops of Tween 80 in 2.5 mL of 7.5% dimethylsulphoxide in PBS to a concentration of 8 μg/μl. Intracranial infusions were conducted using 28G cannulae connected to an infusion pump by polyethylene tubing. 1.0 μl/side was infused into the DH and 0.5 μl/side was infused into the BLA.

### Behavioural equipment

The conditioning chambers (MedAssociates, VT) consisted of two identical illuminated boxes (25 cm × 32 cm × 25.5 cm), placed within sound-attenuating chambers. The box walls were constructed of steel, except by the ceiling and front wall, which were made of perspex. The grid floor consisted of 19 stainless steel rods (4.8 mm diameter; 1.6 mm centre-to-centre), connected to a shock generator and scrambler (MedAssociates, VT). Infrared video cameras were mounted on the ceiling of the chambers (Viewpoint Life Sciences, France) and used to record and quantify freezing behaviour automatically.

### Behavioural Procedures

Rats were conditioned and tested in pairs using previously-established behavioural parameters (21). were subjected to 2 unsignalled footshocks (0.7 mA, 1.5-s), delivered 180 s and 211.5 s into a 273-s session. Two days later, they were returned to the conditioning chamber for a 5-min reactivation session. Rats were infused with SR141716A or vehicle into the DH or BLA either immediately before or immediately after reactivation. They were also injected i.p. with MK-801 or saline, either immediately after the reactivation session (for the pre-reactivation infusion condition) or immediately after the post-reactivation infusion. 3 hours after reactivation, they were returned to the conditioning chamber for a 2-min post-reactivation short-term memory (PR-STM) test. A further 2-min test was conducted 24 hr after reactivation (post-reactivation long-term memory; PR-LTM). All test sessions were video-recorded and automatically quantified for freezing behaviour using video tracking software (Viewpoint Life Sciences, France).

### Statistical analyses

% of time freezing during the test sessions was analysed with repeated measures 4-way ANOVA in JASP 0.8.5.1 (JASP Team 2016) with Timing (pre-vs post-reactivation infusion), Infusion (vehicle vs SR141716A), Injection (Saline vs MK-801), and Test (PR-STM vs PR-LTM) as factors. Planned comparisons analysed the effects of Timing, Infusion and Injection at each test using a 3-way ANOVA. Further planned analyses of simple main effects proceeded by focussing on the effect of Injection, with Timing and Infusion as moderators, to allow an analysis of whether MK-801 had amnestic effects under each sub-condition.

## Results

### Dorsal hippocampus

Figure 1 shows the results obtained with SR141716A infusions in the hippocampus. In the vehicle-infused groups, systemic injections of MK-801 led to a decrease in freezing in comparison to saline-injected controls in PR-LTM tests performed 24 h after reactivation, while no differences were observed in PR-STM tests. SR141716A infusion in the hippocampus abolished the MK-801 effect when performed after reactivation, but had no effect when performed before it.

**Figure 1.**
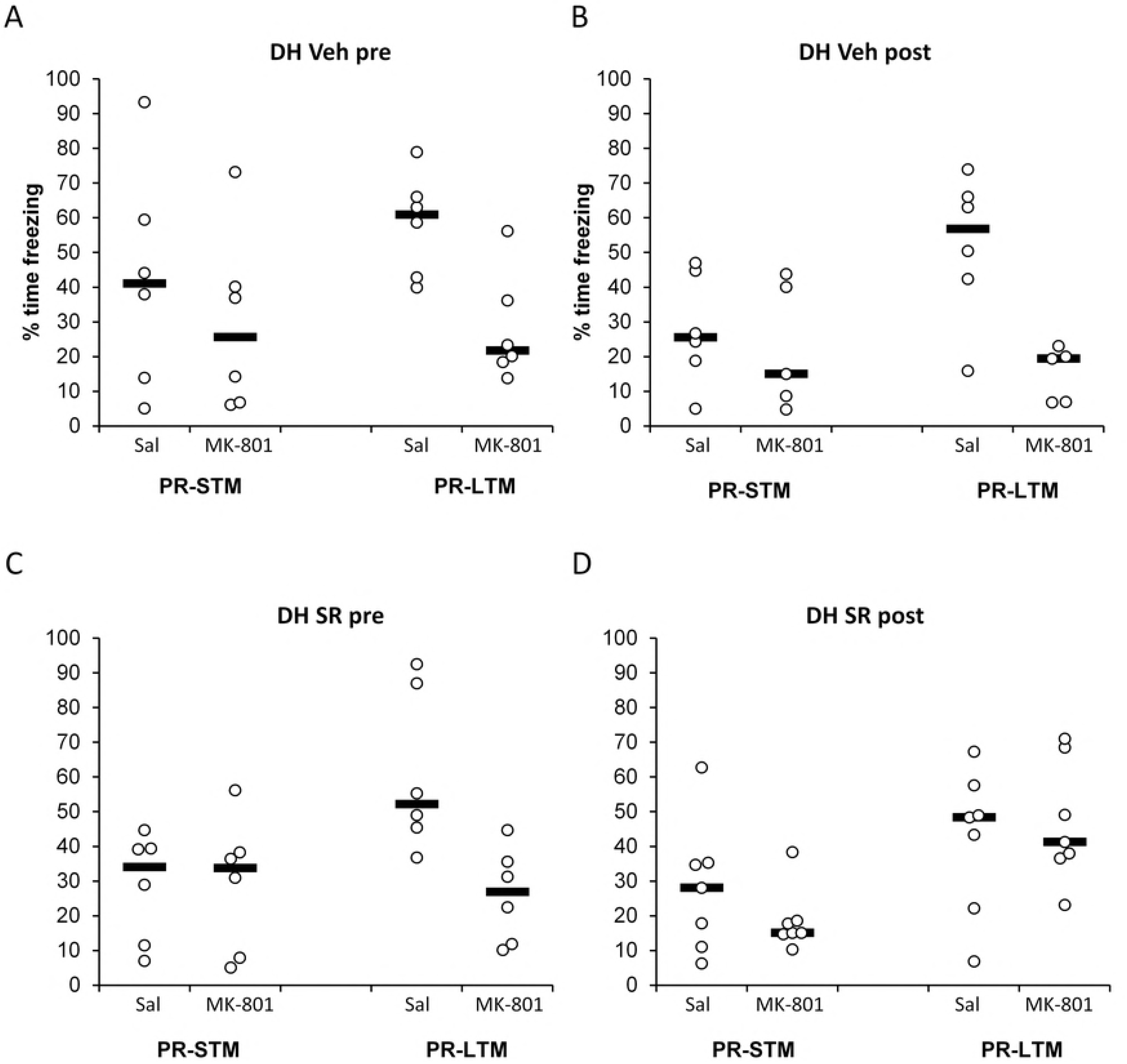
CB1 receptor antagonism in the dorsal hippocampus blocks memory destabilization immediately after, but not before memory reactivation. MK-801 injection (0.1 mg/ml, i.p.) impaired memory reconsolidation in the long-term memory test performed 24h later (PR-LTM) in rats infused with vehicle prior to (A) or immediately after (B) memory reactivation. While pre-reactivation infusion of SR141716A (8 μg/μl) did not alter the amnestic effect of MK-801 (C), post-reactivation SR141716A prevented MK-801-induced impairment in freezing at PR-LTM (D). Statistical analyses confirmed a selective effect in the post-reactivation SR141716A condition (Timing × Infusion × F(1,41)=5.19, p=0.028, η^2^_p_=0.11). No effect of MK-801 injection or SR141716A infusion was observed in the short-memory test at 3h (PR-STM). Data is presented as mean + SEM. n =s 6 for all pre-reactivation groups, 6 for post-reactivation Saline, 5 for post-reactivation MK-801 and 7 for post-reactivation SR141716 + Saline and SR141716 + MK-801.

Analysis of conditioned freezing at PR-STM and PR-LTM tests revealed that the amnestic effect of post-reactivation MK-801 upon contextual fear memory reconsolidation depended upon the timing of intra-dorsal hippocampus infusion of SR141716A vs vehicle (Timing × Infusion × Injection × Test: F(1,41)=5.16, p=0.028, η^2^_p_=0.11). Planned comparisons showed that, at PR-STM, there was no effect of infusion timing, SR141716A or MK-801 (Timing × Infusion × Injection: F(1,41)=0.57, p=0.46, η^2^_p_=0.01; Timing × Infusion: F(1,41)=0.21, p=0.65, η^2^_p_=0.005; Timing × Injection: F(1,41)=0.013, p=0.91, η^2^_p_=0.000; Infusion × Injection: F(1,41)=0.16, p=0.69, η^2^_p_=0.004; Timing: F(1,41)=1.97, p=0.17, η^2^_p_=0.05; Infusion: F(1,41)=0.59, p=0.45, η^2^_p_=0.01; Injection: F(1,41)=1.33, p=0.26, η^2^_p_=0.03). In contrast, at PR-LTM there was further evidence for a timing-dependent SR141716A modulation of reconsolidation disruption by MK-801 (Timing × Infusion × Injection: F(1,41)=5.19, p=0.028, η^2^_p_=0.11). Planned analyses of simple main effects of Injection, with Timing and Infusion as moderators, confirmed that there were impairments in freezing in MK-801-injected rats in both pre-reactivation infusion groups (p’s<0.006) and in the post-reactivation vehicle infusion group (p=0.001), but not in the post-reactivation SR141716A group (P=0.62). Therefore, SR141716A protected against the MK-801-induced impairment of PR-LTM only when infused immediately after the reactivation session.

### Basolateral amygdala

Figure 2 shows the results obtained with SR141716A infusions in the amygdala. Once more, systemic injections of MK-801 led to decreased freezing in PR-LTM tests in both vehicle-infused groups. Unlike in the hippocampus, however, SR141716A attenuated this effect when infused pre-reactivation, and had no effect when infused after it. Once again, no differences were observed in PR-STM tests.

**Figure 2.**
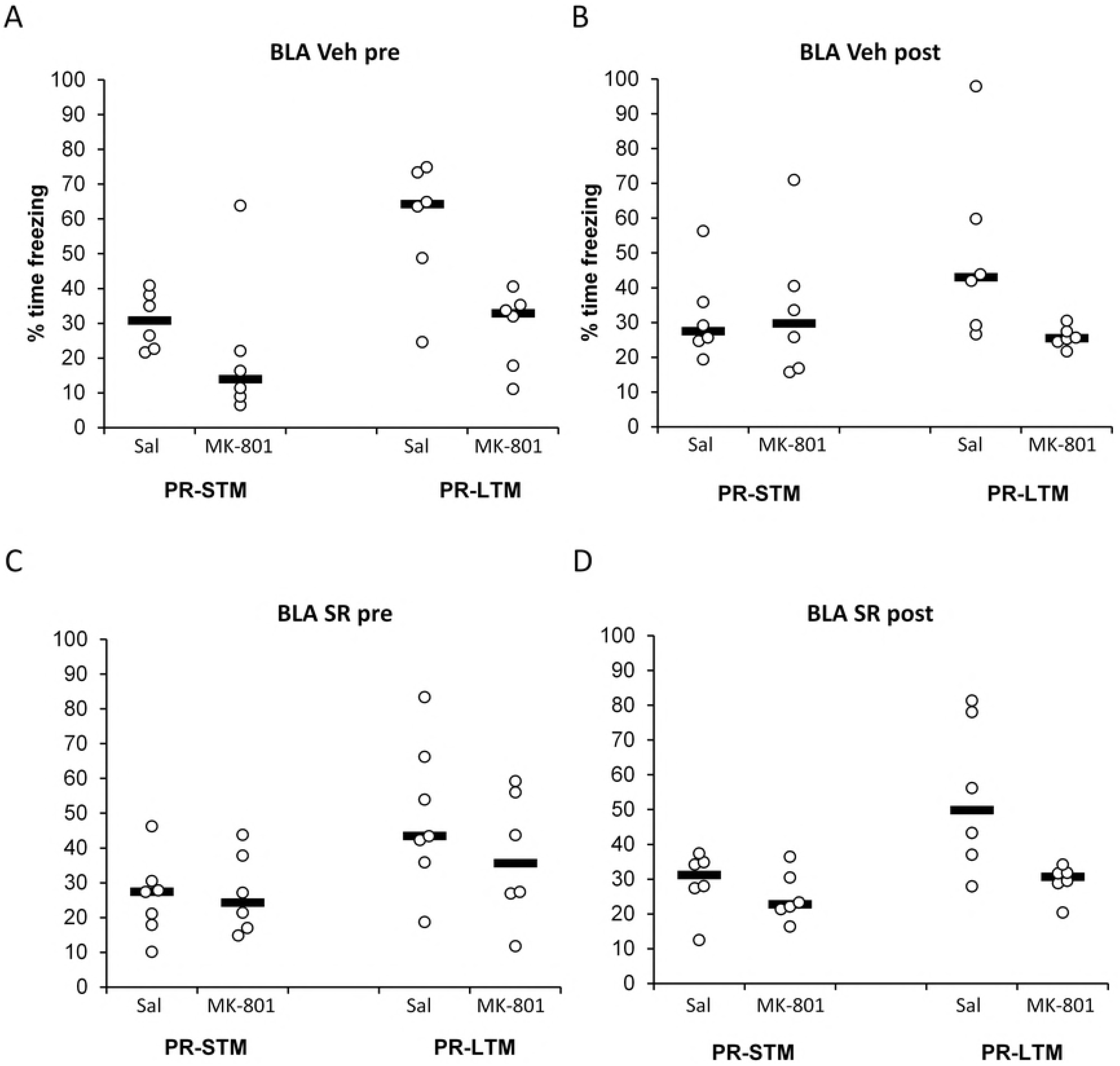
CB1 receptor antagonism in the basolateral amygdala impairs memory destabilization when performed before memory reactivation. MK-801 injection (0.1 mg/ml, i.p.) impaired memory reconsolidation in the long-term memory test (PR-LTM) in the groups infused with vehicle either before (A; simple main effect of injection, p=0.006) or immediately after (B; p=0.023) memory reactivation. While pre-reactivation infusion of SR141716A (8 μg/μl) seemed to prevent the amnestic effect of MK-801 (C; p=0.25), post-reactivation infusion did not (D; p=0.021. However, there was no significant interaction between the timing of infusion and the effect of MK-801 (Timing × Infusion × Injection: F(1,41)=1.11, p=0.30, η^2^_p_=0.026). No effect of MK-801 injection or SR141716A infusion is observed in the short-memory test at 3h (PR-STM). Data presented as mean + SEM. n = 6 per group, except for pre-reactivation SR141716 + Saline, in which n = 7.

Analysis of conditioned freezing at PR-STM and PR-LTM tests revealed that the amnestic effect of post-reactivation MK-801 upon contextual fear memory reconsolidation was not obviously dependent upon the timing of SR141716A infusion (Timing × Infusion × Injection × Test: F(1,41)=0.009, p=0.93, η^2^_p_=0.00; Infusion × Injection × Test: F(1,41)=0.32, p=0.58, η^2^_p_=0.008; Injection × Test: F(1,41)=11.04, p=0.002, η^2^_p_=0.21). Planned comparisons showed that at PR-STM, there was no effect of infusion timing, SR141716A or MK-801 (Timing × Infusion × Injection: F(1,41)=1.11, p=0.30, η^2^_p_=0.03; Timing × Infusion: F(1,41)=0.60, p=0.44, η^2^_p_=0.01; Timing × Injection: F(1,41)=0.15, p=0.70, η^2^_p_=0.004; Infusion × Injection: F(1,41)=0.075, p=0.79, η^2^_p_=0.002; Timing: F(1,41)=0.88, p=0.35, η^2^_p_=0.02; Infusion: F(1,41)=0.50, p=0.49, η^2^_p_=0.02; Injection: F(1,41)=0.42, p=0.52, η^2^_p_=0.02). In contrast, at PR-LTM there was further evidence for an amnestic effect of MK-801 (Injection: F(1,41)=19.77, p<0.001, η^2^_p_=0.33), although this was not clearly dependent upon SR141716A infusion or timing (Infusion × Injection: F(1,41)=0.78, p=0.38, η^2^_p_=0.002; Timing × Infusion × Injection: F(1,41)=1.11, p=0.30, η^2^_p_=0.026). However, planned analyses of simple main effects suggested impairments in freezing in MK-801-injected rats in both post-reactivation infusion groups (p’s<0.024) and the pre-reactivation vehicle infusion group (p=0.006), but not in the pre-reactivation SR141716A group (p=0.25). Therefore, SR141716A appeared to protect against the MK-801-induced impairment of PR-LTM when infused immediately before the reactivation session, although this dissociation was not as clear-cut as that observed in the hippocampus.

## Discussion

Our results show that post-reactivation systemic MK-801 injection disrupted subsequent contextual fear memory expression. However, this memory disruption was seemingly prevented when the CB1 receptor antagonist SR141716A was infused into the BLA or DH. Importantly, SR141716A-mediated protection against amnesia had different temporal windows of efficacy depending upon the locus of infusion. Pre-reactivation infusions were only effective when targeted into the BLA, whereas only intra-dorsal hippocampus post-reactivation infusions prevented MK-801-induced amnesia.

The disruption of contextual fear memory by MK-801 likely reflects an impairment of memory reconsolidation. We and others have previously demonstrated that post-reactivation treatment with NMDA receptor antagonists disrupt reconsolidation of various types of memory (12, 26–28). The post-reactivation timepoint of drug treatment avoids acute effects on the reactivation session itself, and the preservation of contextual fear memory expression at the 3-hr post-reactivation PR-STM test, as expected for reconsolidation blockade (4), rules out non-specific chronic effects of MK-801. While we did not include an operational non-reactivation control condition, the SR141716A-induced protection against the amnestic effect of MK-801 shows that this effect is destabilisation-dependent, and thus likely dependent on reactivation. An alternative account of post-reactivation amnesia focussing on memory integration has been recently proposed (29). While such an account might explain the amnestic effect of MK-801 alone, it is not clear how it would explain the observation that additional treatment with local SR141716A, within specific differential time windows, reverses MK-801-induced amnesia.

The prevention of MK-801-induced reconsolidation disruption by post-reactivation dorsal hippocampal SR141716A replicates a previous study in mice that used intra-hippocampal anisomycin as the amnestic agent (20). This pattern of results, with no effect of SR141716A on its own on the contextual fear memory (and no enhancement of contextual freezing that might offset the disruptive effect of MK-801) has been interpreted as an impairment of memory destabilisation (6, 15, 20, 30), and is consistent with our recent observation that pharmacological agonism of hippocampal CB1 receptors can stimulate the destabilisation of contextual fear memories (22).

While CB1 receptors in the BLA have been studied relatively extensively in fear memory and its extinction, the evidence is more limited when considering memory destabilisation/reconsolidation of contextual fear memories. The CB1 receptor antagonist AM251 had no effect on reconsolidation by itself, but prevented the enhancement of reconsolidation by CB1 receptor agonism in a fear-potentiated startle setting (31). This was interpreted as a purely pharmacological effect (i.e. the AM251-blockade of CB1 receptors directly preventing pharmacological agonism), although it is not inconsistent with a potential effect of CB1 receptor antagonism in preventing memory destabilisation.

However, Ratano et al (32) observed that AM251, at a substantially lower dose (300 ng/side c.f. 20 μg/side), did impair post-reactivation long-term memory. This effect was shown with AM251 infusions immediately after, but not 30 min prior to, memory reactivation and appeared to be mediated by the dysregulation of GABAergic signalling in the BLA (32). This contrasts with our observation that SR141716A had no effect alone when infused immediately before or after reactivation. It remains unclear what accounts for this discrepancy, and what are its implications for our interpretation of an intra-BLA SR141716A-mediated impairment of contextual fear memory destabilisation. A difference in memory type (cued vs contextual fear) exists, but may not be important, given that the BLA is hypothesised to have a conceptually similar role in both settings, associating the CS or contextual representation with the US (33–35). Another difference is the CB1 antagonist employed: while AM251 and SR141716A are structurally similar, and both have “inverse cannabimimetic effects” consistent with pharmacological inverse agonism (36), there is some evidence that the two drugs may differ in their affinity for an unidentified central vanilloid VR1-like receptor (37).

Our BLA SR141716A results are not only potentially inconsistent with those of Ratano et al (32), but less clear-cut statistically than our DH SR141716A results. The conclusion that intra-BLA SR141716A disrupts contextual fear memory destabilisation only when infused prior to memory reactivation results from planed analyses of simple main effects, and should be considered as preliminary in the absence of conclusive evidence for a dependence of MK-801 effects upon either the timing of the BLA infusion or its content (SR141716A vs vehicle). Nevertheless, there is a clear difference between the BLA and DH results, with stronger evidence for intra-dorsal hippocampal SR141716A having destabilisation-impairing effects only when infused after reactivation. This temporal pattern is largely consistent with previous studies using different destabilisation-inhibitors, with pre-reactivation selectivity in the BLA (3) and post-reactivation sufficiency in the hippocampus (20, 36). Our study adds to this picture the fact that, in the hippocampus, the effects of destabilisation blockade seem to be restricted to the post-reactivation period. We also demonstrate different temporal windows of efficacy of destabilisation blockade between the amygdala and hippocampus when the same memory task, amnestic agent and destabilisation inhibitor are used. Therefore, the differential effects are not likely to be due to pharmacodynamics or pharmacokinetics of different drugs, nor to potential differences in the engagement of destabilisation mechanisms because of discrepant memory settings and strength of conditioning.

This opens up an interesting question concerning whether memory destabilisation during reconsolidation represents a single, unified phenomenon, or whether distinct phenomena involving CB1 receptors in different structures lead to a similar behavioural outcome of memory weakening when reconsolidation is blocked. As destabilisation shares common molecular mechanisms across brain structures (3, 16, 38), it is possible that the different temporal profiles of CB1 involvement reflect the distinct temporal dynamics of each structure’s role in fear conditioning. In this view, destabilisation mechanisms are engaged at a later stage in the hippocampus, but are involved in plasticity processes that are similar to those in the amygdala, as suggested by the common behavioural outcome of destabilisation blockade in both structures.

A second possibility, however, is that the mechanism through which CB1 receptor activation leads to memory destabilisation is different in both structures. Although the pharmacological profile of destabilisation in the amygdala and in the hippocampus is generally similar, some of the molecular mechanisms involved – which include CB1 receptors, the ubiquitin-proteasome system (15, 20) and AMPA receptor endocytosis (9, 17) in both structures, L-type voltage-gated calcium channels in the hippocampus (20) and calcineurin in the amygdala (11) – are also involved in other forms of behavioural and synaptic plasticity, such as memory extinction (1), normal forgetting (39) and homeostatic synaptic downscaling (40, 41). Thus, it is possible that the CB1 receptor might be part of a more general plasticity system that is engaged in the amygdala and the hippocampus for different purposes during memory updating.

Finally, another open question is how other functions of the CB1 receptor in the amygdala, such as the mediation of different forms of memory extinction (42, 43) and acute fear relief (44) relate to its role in memory destabilisation during contextual re-exposure. It is interesting to note that CB1 receptors seem to be particularly important for within-session freezing decrease (45), a phenomenon that can temporally co-occur with memory labilization during re-exposure. Although no correlation has been shown between the degree of freezing decrease during re-exposure and the effect of post-reactivation injections of MK-801 (21), it is nevertheless possible that the decrease in fear during re-exposure mediated by CB1 could play a role in setting off labilization mechanisms, which might later mediate memory updating in the hippocampus and other structures.

These and other matters, however, remain open to further studies, which might reveal whether the temporal dissociation between labilization in the hippocampus and amygdala observed with CB1 receptors occurs with other molecular targets as well. In the meantime, our results show that memory destabilisation during reconsolidation of contextual fear conditioning is a complex process that cannot be pinpointed to a single structure or time point, and that the temporal dynamics of the engagement of destabilisation mechanisms may differ between brain structures.

## Acknowledgments

The authors thank David Barber for technical support. The research was conducted with funding from the UK BBSRC (grant number BB/J014982/1), FAPERJ (grant number E-26/010.002674/2014) and the University of Birmingham.

